# A type II Thoeris anti-phage defense enacts growth arrest independent of NAD depletion

**DOI:** 10.64898/2026.01.22.701046

**Authors:** Frank Englert, Oleg Dmytrenko, Chase L. Beisel

**Affiliations:** Helmholtz Institute for RNA-based Infection Research (HIRI), Helmholtz-Centre for Infection Research (HZI), Würzburg, Germany; Medical Faculty, University of Würzburg, Würzburg, Germany

## Abstract

Thoeris immune systems protect bacteria against invading phages through production of a signaling molecule by ThsB that activates the effector ThsA. Across the four defined types, the two most dominant (I and II) have been associated with abortive infection, with type I acting through the depletion of the essential coenzyme NAD^+^. Here, we show that a previously uncharacterized type II Thoeris system from *Escherichia coli* ATCC 25922 deviates from this paradigm. This system consists of the effector protein ThsA and two distinct signaling proteins ThsB1 and ThsB2. Heterologous expression of *thsA* and *thsB2* confers anti-phage defense, while *thsB1* is cytotoxic when expressed in common *E. coli* lab strains. Furthermore, while phage infection drives growth arrest, we could not detect any measurable decrease in NAD^+^ levels as well as standard markers of cell death. Together, these results suggest that Thoeris contains even greater functional diversity within the defined system types.

## INTRODUCTION

Bacterial immune systems collectively comprise a wide range of defense mechanisms employed by bacteria to protect against bacteriophage (phage) infection (1). These include various innate immune systems, restriction-modification systems being the most distributed, as well as the adaptive immune system CRISPR-Cas (2). Systematic bioinformatic searches have uncovered more than 190 bacterial defense systems, with over 60 experimentally characterized and more than 750 additional unique proteins predicted to function in antiphage defense (2–11).

One of these defense systems, called Thoeris, was discovered in 2018 and identified in 4% of bacterial genomes, representing one of the most prevalent forms of bacterial immunity (3). It consists of a protein-encoding gene named *thsA* paired with one or more protein-encoding genes named *thsB*. ThsB senses the phage infection, producing a signaling molecule derived from nicotinamide adenine dinucleotide (NAD^+^) through its Toll-interleukin receptor (TIR) domain. The diffusible signaling molecule then binds to ThsA to activate its effector function, which blocks further propagation of the phage.

Four types of Thoeris systems have been defined to-date principally based on the annotated domains within ThsA (**Fig. 1a**). Type I ThsA contains a sirtuin (SIR2)-like domain and a SMF/DprA-LOG (SLOG) domain (12), type II ThsA contains a a Macro domain and putative trans-membrane domains (12), type III ThsA contains a SLOG domain (12), and type IV ThsA contains a Caspase domain (13). Beyond ThsA, other identified components and features differentiate the types. For instance, ThsB proteins show large diversity across the types, with 11 sub-families defined for those associated with types I and II systems (12). Separately, the identified signaling molecules synthesized by ThsB and triggering ThsA vary between types, with type I systems utilizing isomers of a cyclic adenosine diphosphate ribose (cADPR) with the O-glycosidic bond formed at the 1′, 2′ or 3′ position (14–16), type II systems utilizing a histidine-linked adenosine diphosphate ribose (17) and imidazole adenine dinucleotide (12), and type IV systems utilizing N7-cADPR (13). Finally, the modes of immune defense enacted by ThsA vary across types, albeit with a common tendency to enact abortive infection defined in part by immunity-driven cell death. In particular, representative systems from types I, II, and IV were shown to drive abortive infection respectively through cleavage of NAD^+^, an essential coenzyme (18, 19), an unknown mechanism that has been linked to ThsA’s transmembrane domains (12), or indiscriminately degrading cellular proteins (13). In contrast, type III systems lack an abortive infection phenotype, where its activity is hypothesized to depend on its NucS nuclease effector domain (6). This current knowledge underscores a range of mechanisms and functions within the architecture of Thoeris systems and suggests that greater diversity remains to be revealed.

**Figure 1.**
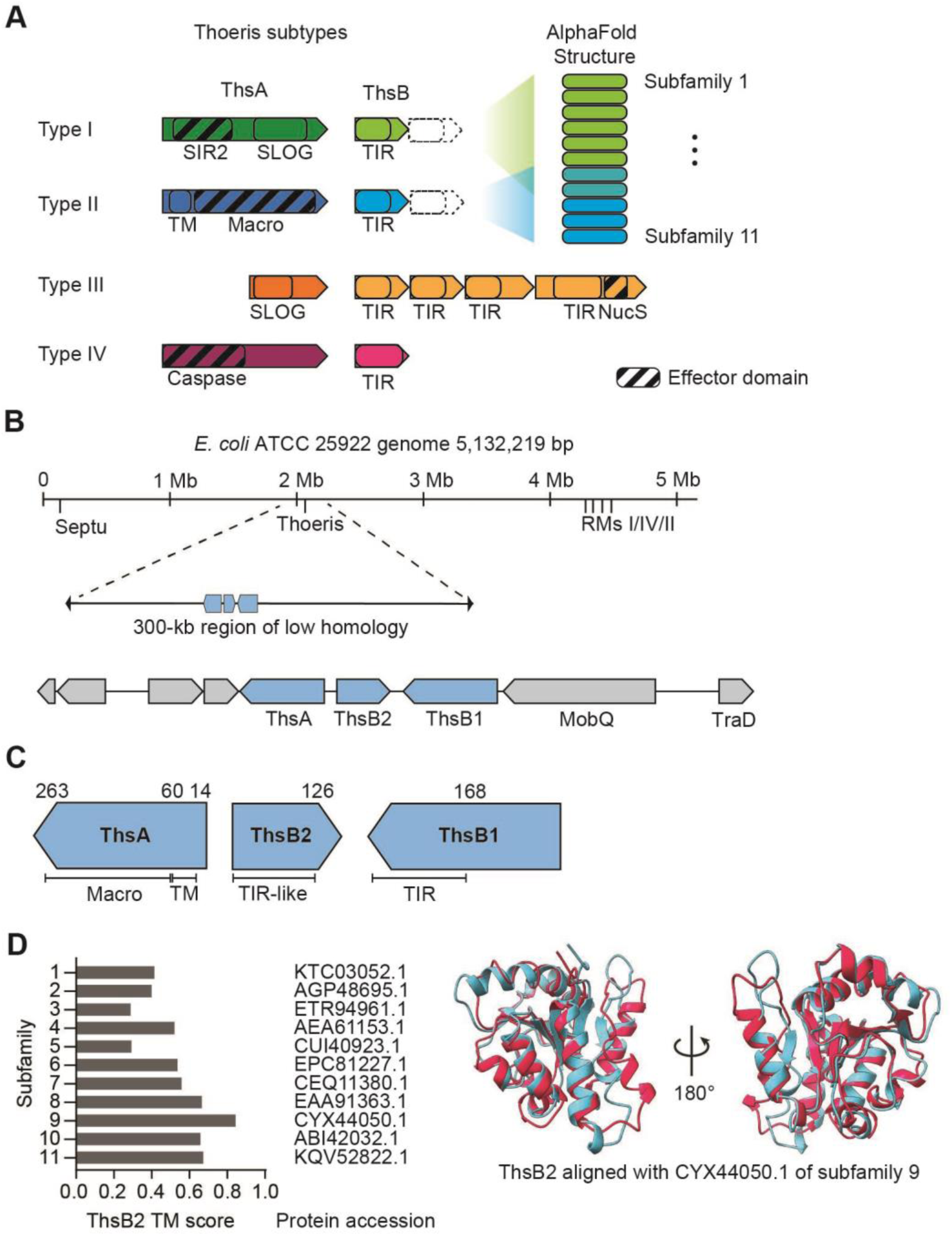
*E. coli* ATCC 25922 encodes a type II subgroup 9 Thoeris system in a three-gene locus. **A**: Overview of the different types of discovered Thoeris systems. Previously characterized Thoeris systems are grouped into four distinct types, with type I and type II further distinguished into 11 subfamilies. All types have in common one or more TIR-domain containing genes (called *thsB* in type I and II, *thcB* in type III and *PsTIR* in type IV). The categorization is based on the effector domain (highlighted with diagonal bars), which is unique for every type. **B**: Genome context of the ATCC 25922 type II Thoeris system: An overview of the 5, 132, 219 bp genome of *E. coli* ATCC 25922, with the locations of known phage-defense systems highlighted. The Thoeris system, highlighted in blue, is located in a part of the genome that is not homologous to other *E. coli* strains, but is not in proximity to other known phage defense systems. **C**: Domain annotation: The Thoeris locus, consisting of *thsA*, *thsB2* and *thsB1*, with their predicted functional domains annotated. The exact amino acid boundaries are shown on top. *ThsB2* is predicted to contain a TIR-like domain (pf08937), similar to other Thoeris systems, while *thsB1* is predicted to contain a TIR domain (pf13676). **D**: Alignment of the predicted structures of the ATCC 25922 ThsB2 protein (red) with the *Streptococcus suis* ThsB (blue; CYX44050. 1). CYX44050. 1 was chosen as representative of the subgroup 9 of type II Thoeris systems. TM-align of the two structures gave a TM-score of 0. 84431, where 1 would indicate a perfect match and < 0. 2 would correspond to randomly chosen unrelated proteins.

In this study, we characterized a type II Thoeris system native to the *E. coli* strain ATCC 25922. We found that this system prevents phage infection but without exhibiting an abortive infection phenotype, Furthermore, we show that cellular NAD^+^ levels remain unchanged following phage infection and rule out other common mechanisms associated with membrane disruption and abortive infection. These findings contrast with the previously characterized type II Thoeris systems and underscore the functional diversity of Thoeris-mediated defense.

## RESULTS

### *E. coli* ATCC 25922 encodes a type II Thoeris system

To investigate the diversity of type II Thoeris systems, we focused on the *E. coli* strain ATCC 25922 and its native Thoeris system, which was identified as type II based on the Macro domain identified in *thsA* (12). This system is located in a 300-kb region of the genome that shares little homology to other laboratory strains of *E. coli* based on progressive MAUVE alignment (**Figs. 1B and S1**) (20). The Thoeris locus is separated by at least 1 Mb from the other annotated anti-phage defense systems (Septu, Lamassu-Mrr, PsyrTA, SanaTA, type I, II, and IV restriction-modification systems) and is flanked upstream only by genes encoding putative proteins and downstream by conjugation genes *mobQ* and *traI*. The Thoeris locus consists of one *thsA* gene and two *thsB* genes (*thsB1*, *thsB2*), with only *thsB2* oriented in the opposite direction. Similar to previously described type II Thoeris systems, *thsA* contains a Macro domain and a putative trans-membrane (TM) domain (**Fig. 1C**) (3, 12), while *thsB1* contains a TIR-domain (pf13676) and *thsB2* contains a DUF1863 (pf08937) domain described as a TIR-like domain (21, 22).

To determine the subgroups in which ThsB1 and ThsB2 belong (12), we used Alphafold3 (23) to predict the structures of ThsB1 and ThsB2 and compared them to representative ThsB structures using TM-align (24). For ThsB1, only the conserved TIR-domain aligned with representative subgroup structures, with TM-scores below 0. 5, indicating minimal structural similarity (**Table S1)**. The N-terminal part of ThsB1 lacked other predicted domains, and a DALI search did not reveal any meaningful alignments (25). In contrast, ThsB2 showed strong structural similarity to ThsB CYX44050. 1 (**Fig. 1D, Table S1**), which is associated with subgroup 9.

### *thsA* and *thsB2* in *E. coli* ATCC 25922 protect against phage infection

To assess the extent to which the Thoeris system in *E. coli* ATCC 25922 protects against phage infections under native conditions, we deleted the *thsAB2B1* locus and exposed the deletion strain to 73 diverse *E. coli* bacteriophages (**Fig. 2A**) (26). A majority of the phages failed to form plaques, regardless of infecting the wild-type or deletion strain. The phages that were able to form plaques showed no difference in plaquing efficiency or plaque morphology between wild-type and deletion strains, indicating that deleting the native *thsAB2B1* locus did not enhance sensitivity to phage infection under the tested conditions.

**Figure 2.**
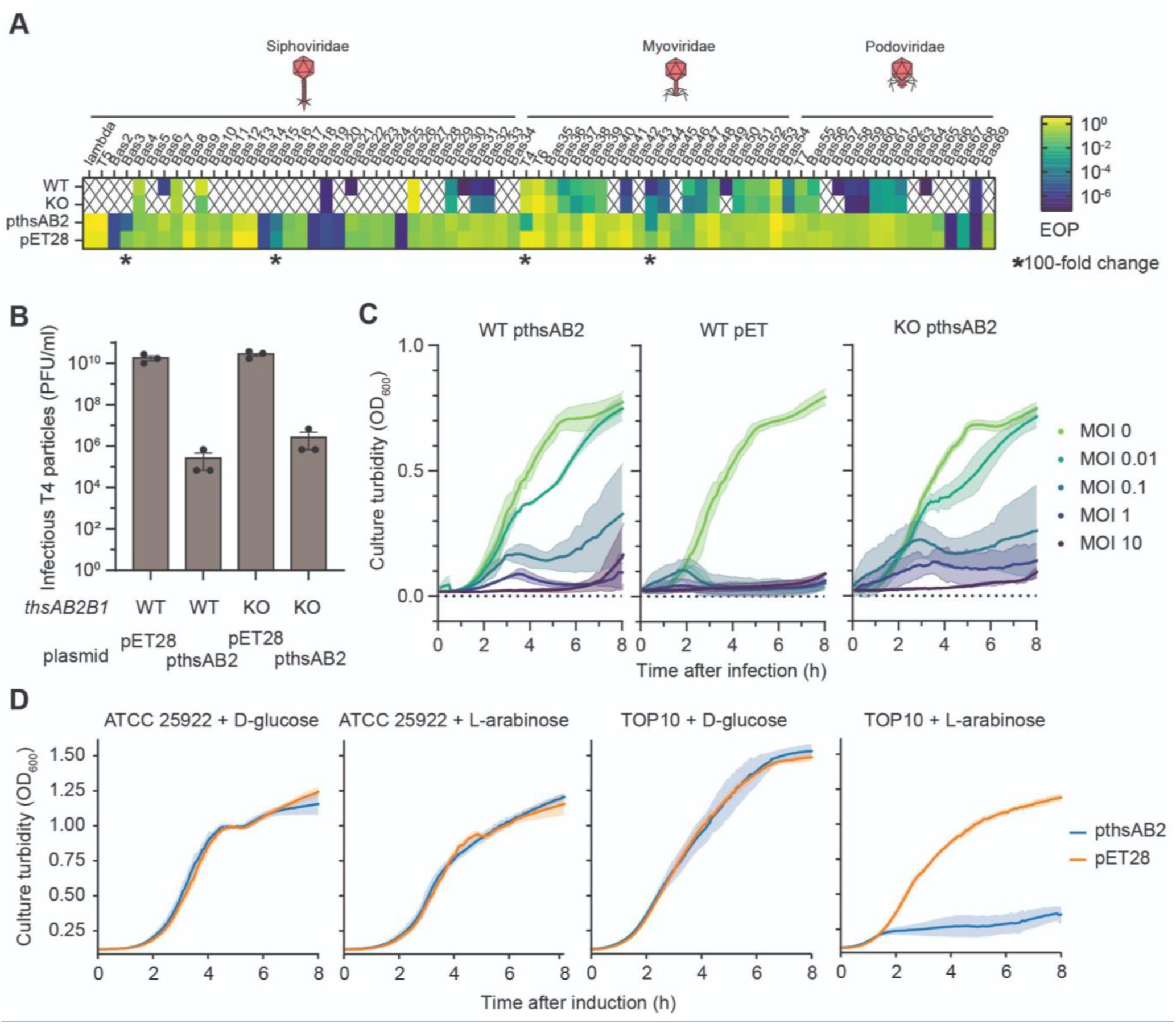
The ATCC 25922 *thsAB2* locus protects against various phages when heterologously expressed. **A:** *E. coli* ATCC 25922 is resistant to most of the Basel phage collection, even in the absence of Thoeris. Number of phage plaques forming on the native ATCC 25922 host (WT), with the *thsAB2B1* locus knocked out (KO) or on a susceptible host carrying an empty vector plasmid (pET28) or the same vector carrying the *thsAB2* locus (pthsAB2). An asterisk highlights the phages showing >100-fold difference between pthsAB2 and pET28. A crossed out square means no plaques detected. Each square represents the average of two biological replicates. EOP: efficiency of plating. **B:** Susceptibility of *E. coli* ATCC 25922 to phage T4 infection. The *E. coli* strain contains the native Thoeris *thsAB2B1* locus or expresses the *thsAB2* locus from a plasmid, compared to the *thsAB2B1* knockout strain and an empty vector control. **C:** Growth curves of ATCC 25922 WT and *thsAB2B1-*deletion strains, carrying *thsAB2* plasmid, infected by phage T4 at varying MOIs. ATCC 25922 WT carrying an empty vector control is shown as negative control, being susceptible to phage T4. **D:** Heterologous expression of the *thsB1B2A* locus, with *thsB1* under the control of an arabinose inducible promoter. Growth curves of *E. coli* MG1655 and ATCC 25922 strains heterologously expressing *thsAB2B1* under inducing (+ L-arabinose) or repressing (+ D-glucose) conditions. Lines and shaded regions represent the mean and standard deviation of three independent cultures.

We therefore ported the Thoeris system into a heterologous expression system by cloning the *thsAB2B1* locus, including native promoters, into a multicopy plasmid. We were unable to clone the full *thsAB2B1* locus but successfully cloned the contiguous *thsAB2* region. To determine if this shortened locus could confer protection against phage infection, the pthsAB2 plasmid was transformed into the susceptible *E. coli* MG1655 strain and exposed to the same library of phages (**Fig. 2A**). Heterologous expression of *thsAB2* conferred resistance against several phages, with phage T4, Bas3, Bas15 and Bas43, yielding a 100 -1, 000-fold reduction in plaquing compared to an empty vector control. The resisted phages belong to distinct morphotypes (Syphovirus, Myovirus), families (Drexlerviridae, Straboviridae, unassigned) and recognize different terminal receptors (FhuA, TsX, OmpC), implicating no specific mechanism of activation.

To determine whether plasmid-based expression of *thsAB2* also confers protection in the native host, we introduced the same pthsAB2 expression plasmid into both the wild-type (WT) ATCC 25922 strain and Δ*thsAB2B1* mutant. We tested susceptibility to phage T4, chosen because it efficiently formed plaques on the WT and deletion strains (**Fig. 2A**). *thsAB2* expression resulted in a 105-fold reduction in plaque formation in the WT and the Δ*thsAB2B1* strains, comparable to the protection observed when expressing *thsAB2* in *E. coli* MG1655 (**Fig. 2B**). We further monitored infection dynamics in liquid culture at multiplicities of infection (MOI) ranging from 10 to 0. 01 (**Fig. 2C**). Across all MOIs, *thsAB2*-expressing cultures maintained higher cell densities than empty-vector controls, although growth was increasingly suppressed at higher phage loads. Together, these results demonstrate that plasmid-based expression of the *thsAB2* locus in its original strain and a heterologous strain protect against diverse phages.

### Induced plasmid expression of *thsB1* is toxic in heterologous *E. coli*

As we were only able to clone the Thoeris locus without *thsB1*, we placed *thsB1* under an L-arabinose-inducible promoter with a modified 5′ untranslated region, while *thsA* and *thsB2* remained under their native promoter. Under non-inducing conditions, bacterial growth in liquid culture matched that of an empty-vector control. However, induction of *thsB1* expression blocked growth in *E. coli* TOP10 as well as in *E. coli* BL21(DE3) (**Figs. 2D and S2a**). Interestingly, no growth defect was observed in the native Δ*thsAB2B1* strain (**Fig. 2D**), suggesting that the native strain lacks sensitivity or blocks cytotoxicity. The cytotoxicity did not appear to be linked to ThsB1’s anticipated ability to synthesize NAD-derived signaling molecules, as mutating any of the three WxxxE motifs conserved among TIR domains (27–29) did not restore growth under inducing conditions **(Fig. S2b)**. Whereas these findings indicate that overexpression of *thsB1* is toxic in the non-native *E. coli* strains, this toxicity appears to be independent of any NADase activity.

### Conserved features of ThsA and ThsB2 are essential for phage defense

To further investigate the *thsA-thsB2* locus, we assessed whether the both genes and their conserved domains are essential for anti-phage defense. Consistent with previously characterized Thoeris systems (3), expression of either ThsA or ThsB2 alone failed to confer protection against phage infection (**Fig. 3B**). Given the predicted N-terminal transmembrane helices of ThsA, we examined their importance for function. Truncation of *thsA* by removal of either the first or both helices abolished phage defense (**Fig. 3A-B**). Similarly, mutation of the conserved tryptophan within the single WxxxE motif of ThsB2 completely eliminated defense activity (**Fig. 3B**). Thus, both ThsA and ThsB2 as well as their conserved transmembrane and TIR domains are essential for anti-phage defense.

**Figure 3.**
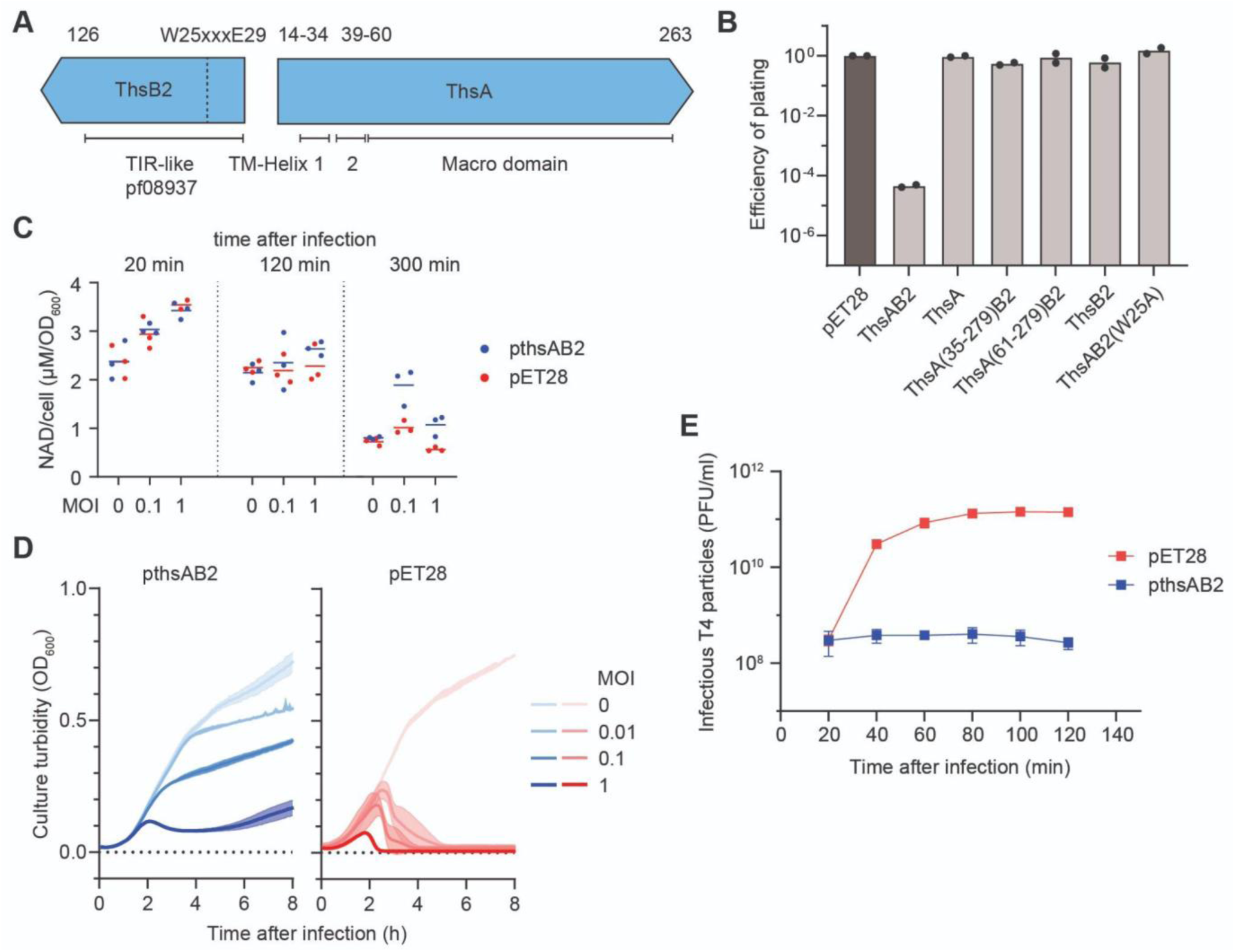
*thsAB2* is sufficient to confer phage resistance when heterologously expressed. **A:** Overview of the truncated Thoeris locus, consisting only of *thsB2* and *thsA*. The native sequence upstream of *thsB2* and downstream of *thsA* was included for the heterologous expression from a multicopy plasmid. The predicted domains are annotated below. The residue borders and the tested point mutation are numbered above. **B:** Efficiencies of plaquing of phage T4 against *E. coli* heterologously expressing the Thoeris system or an empty vector control, compared to a susceptible *E. coli* host. Thoeris was expressed in the form of the truncated *thsAB2* locus, as well as *thsA* and *thsB2* alone. Functional mutations of the thsAB2 locus were tested, by truncating the *thsA* transmembrane domains or mutating the *thsB2* WxxxE motif. **C:** NAD assay to determine the NAD concentration of Thoeris protected *E. coli* infected by phage T4 at varying MOIs. Cultures were harvested at three timepoints post infection, representing an immediate reaction to infection (20 min), tolerance of several infection cycles (120 min) and a timepoint at which unprotected cultures start to collapse (300 min). Each assay was done in triplicate and the calculated NAD concentration was normalized to OD = 1. **D:** Growth curves of *E. coli* heterologously expressing *thsAB2* (pthsAB2) or an empty vector control (pET28), after infection with phage T4 at different MOIs. Each curve shows the average of three independent replicates, with the shaded area indicating the standard deviation. **E:** One step growth curves of phage T4 on *E. coli* heterologously expressing *thsAB2* or an empty vector control. Phage titer was determined in 20 min intervals. Each point is the average of three replicates, with the error bars showing the standard deviation.

### *thsAB2*-mediated defense does not deplete cellular NAD

Previously characterized Thoeris type I systems deplete cellular NAD^+^ through the ThsA SIR2 domain (18). The mechanism for type II systems remains unknown (12), although Macro domains have been reported to bind and, in some cases, hydrolyze NAD^+^ (12). To test whether NAD depletion contributes to type II Thoeris defense, we quantified NAD^+^ concentration in *E. coli* heterologously expressing *thsAB2* following infection with phage T4 at varying MOIs (**Fig. 3C**). NAD levels were measured using a colorimetric assay based on a lactate dehydrogenase cycling reaction. Samples were collected at multiple timepoints during the infection cycle (**Fig. S3**), but no significant differences in NAD concentrations were observed. This indicates that this Thoeris system does not involve cellular NAD^+^ depletion during phage defence, suggesting an alternate mode of antiviral activity.

### *thsAB2* prevents propagation of phage T4 without culture collapse

Previously characterized type I, II and IV Thoeris systems exhibit features of abortive infection, typified in part by a premature collapse of the bacterial culture that prevents phage propagation (3, 18, 30). To determine whether *thsAB2* mediated defense follows a similar mechanism, we challenged *E. coli* heterologously expressing *thsAB2* with T4 phage at varying MOIs and compared the growth to that of unprotected controls. Unprotected cultures collapsed within two hours and did not recover. In contrast, *thsAB2*-expressing cultures exhibited growth arrest but plateaued at MOI-dependent densities rather than collapsing (**Fig. 3D**).

To assess whether this phenotype reflects a reduction in viable phages, we quantified phage titers over the course of several infection cycles (**Fig. 3E**). In empty-vector controls, T4 replicated with the expected 30-minute infection cycle (31), resulting in a sharp increase in plaque formation. In contrast, phage titers remained constant in *thsAB2-*expressing cultures throughout the 120 minute observation period. Together, these results demonstrate that *thsAB2* blocks phage propagation through inhibition of cell growth rather than driving culture collapse.

### *thsAB2*-mediated defense does not cause abortive infection

As *thsAB2*-mediated defense did not drive culture collapse, we asked if other hallmarks of cell death indicative of abortive infection could be identified. To monitor cell death during phage infection, we used propidium iodide (PI), an intercalating dye that produces a fluorescent signal upon binding to double stranded DNA released during lysis. In unprotected bacteria exposed to T4 phage, fluorescence increased in parallel with culture collapse, consistent with phage-induced lysis (**Fig. 4A**). In contrast, *thsAB2*-expressing cultures showed only a minor fluorescence increase at the highest tested MOI. At lower MOIs, fluorescence remained unchanged, consistent with the growth curves showing plateauing rather than culture collapse.

**Figure 4.**
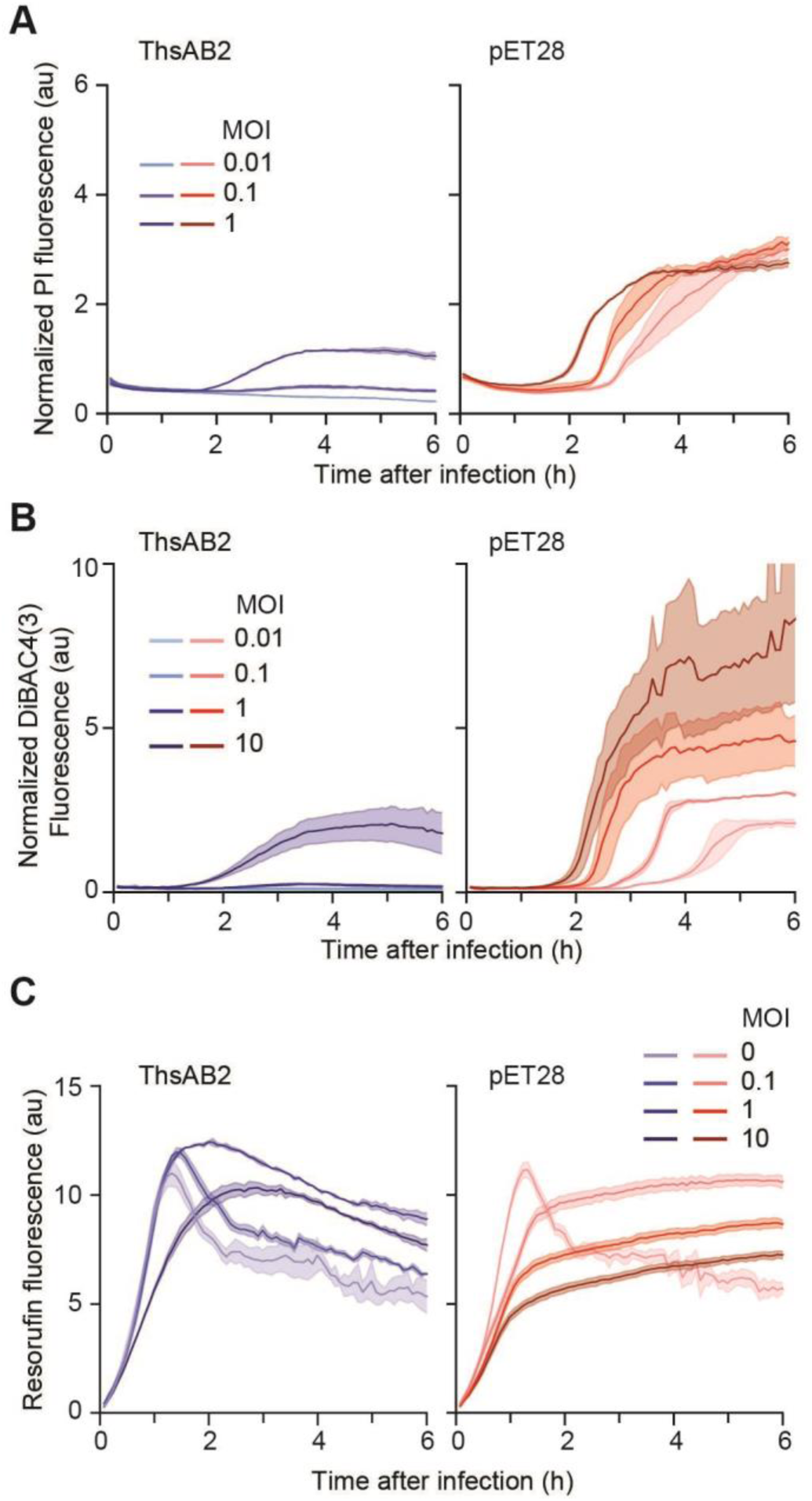
*thsAB2*-mediated phage defense shows no indication of abortive infection. **A:** PI indicator assay to evaluate cell death following phage infection of Thoeris protected (left) and unprotected (right) *E. coli* following phage T4 infection. Each curve shows the average of three independent replicates, with the shaded area indicating the standard deviation. **B:** DiBAC4(3) indicator assay to evaluate membrane polarization of Thoeris protected (left) and unprotected (right) *E. coli* following phage T4 infection. Each curve shows the average of two independent replicates, with the shaded area indicating the standard deviation. **C:** Resazurin indicator assay to compare metabolism rate of Thoeris protected (left) and unprotected (right) *E. coli* following phage T4 infection. Each curve shows the average of two independent replicates, with the shaded area indicating the standard deviation.

As ThsA contains a putative transmembrane domain, we asked whether *thsAB2*-mediated defense involves membrane perturbation. Abortive infection systems can depolarize the membrane, for example, through pore formation, leading to cell death. To test this, we used the membrane potential probe DiBAC4(3), which enters depolarized cells and binds intracellular proteins, increasing the fluorescent intensity. As in the PI assay, an increase in fluorescence was observed only at the highest MOI **(Fig. 4B**), whereas lower MOIs showed no detectable change despite strong growth inhibition. This indicates that the membrane potential stays intact during phage defense.

Since *thsAB2*-expressing cultures displayed growth inhibition but no signs of cell death, we investigated whether bacteria enter a dormant or torpid state characterized by reduced metabolism. To do so, we used resazurin, a weakly fluorescent, cell permeable dye that is reduced to highly fluorescent resorufin in metabolically active cells. Resorufin is further reduced to the non-fluorescent dihydroresorufin, leading to decay of fluorescence over time. In uninfected cultures containing resazurin, the fluorescence of both the *thsAB2-*expressing culture and the empty vector control peaked after 1. 5 hours, defining the baseline metabolic rate (**Fig. 4C**). In infected *thsAB2*-protected cultures containing resazurin, this peak remained unchanged up to MOI 1, and shifted only modestly at MOI 10. This largely unchanged metabolic activity remained even under strong growth inhibition. These observations indicate that reduced metabolism is not responsible for growth arrest during *thsAB2-*mediated defense.

Together with the absence of culture collapse, limited cell death, and absence of membrane depolarization, these results demonstrate that *thsAB2* protects against phage infection without triggering hallmarks of abortive infection.

## DISCUSSION

In this study, we investigated a type II Thoeris from the *E. coli* strain ATCC 25922 system in both its native and heterologous contexts. Although *thsB1* proved toxic when expressed heterologously, *thsA and thsB2* together were sufficient to protect against several phages. We demonstrated that phage propagation was efficiently inhibited without inducing an abortive infection phenotype.

The ATCC 25922 strain is highly resistant against most tested phages. Resistance persisted even when the Thoeris system was deleted, suggesting the presence of additional antiphage mechanisms. ATCC 25922 carries multiple prophages (32), which are known to frequently encode phage defence systems (33–36). Beyond prophages, its genome is nearly 1 Mb larger than that of common laboratory *E. coli* strains and includes two plasmids, either of which could harbor additional defence mechanisms. Using DefenseFinder (2, 5, 37–39), we identified Septu, Lamassu-Mrr, restriction-modification, as well as Psyr and Sana toxin-antitoxin systems that may contribute to the phage resistance of this strain. Morphology could be another source of phage resistance, especially biofilms have been shown to protect susceptible bacteria from phage infection. ATCC 25922 has been shown to be a strong biofilm producer (40) that could also contribute to phage resistance (41), even though the experimental conditions tested in this work would not be expected to promote biofilm formation. Taken together, the extensive phage resistance of ATCC 25922 likely arises from a combination of multiple distinct defense systems and strain-specific physiological traits, underscoring the multifactorial nature of antiviral immunity in this strain.

The co-occurrence of multiple *thsB* genes has previously been linked to an expanded range of recognized phages by the Thoeris system (18), yet the cause of *thsB1*-associated toxicity during heterologous expression remains unclear. Beyond NAD^+^ hydrolysis, ThsB proteins have been shown to catalyze base-exchange reactions with pyridines, pyridine-fused heterocycles, and other five-membered heterocyclic compounds (12). The structural diversity among ThsB subfamilies suggests additional, as-yet-unknown mechanisms of function. It also remains unresolved whether *thsB1* toxicity in the native host is through, for example, epigenetic regulation or by an unidentified antagonist.

Thoeris type II have been hypothesized to act through abortive infection, owing to their putative membrane association and similarity to other Thoeris variants. However, in this study none of the canonical phenotypes of abortive infection — culture collapse, cell death or membrane depolarization — were observed. Instead, we observed that cell density plateaued rather than collapsed, resembling protection conferred by systems that induce growth arrest without killing the host, such as by type VI CRISPR-Cas systems, Viperins, long-A pAgos, or Shango (42–45). Mechanisms shutting down cells while preventing phage replication, such as dormancy or growth arrest would be consistent with our PI staining results, which showed minimal cell death, although the underlying mechanism remains to be elucidated.

Phage infections of bacteria have major industrial consequences, particularly in the dairy and fermentation industries (46). To mitigate these effects, bacterial strains are often equipped with engineered phage defense systems, most notably CRISPR-Cas (47, 48). A recent study showed that combining distinct bacterial immune systems can produce synergistic protection against a broader range of phages (49). In particular, a type I Thoeris system combined with CRISPR-Cas provided greater immunity than either system alone. Our work reveals that Thoeris defenses extend beyond abortive infection and can suppress phage propagation without host lethality. This mechanistic diversity broadens the available toolkit of bacterial immune systems, offering new opportunities for rationally combining diverse defense mechanisms to achieve optimal protection in industrial and synthetic biology contexts.

## METHODS

### Bacterial strains and growth conditions

*E. coli* strains used in this study were cultivated in LB medium (10 g/L Tryptone, 5 g/L NaCl and 5 g/L yeast extract) at 37°C, shaking orbitally at 220 rpm, or on LB agar (Carl Roth, Cat. # X965. 3) plates incubated at 37°C, unless otherwise stated. To make soft agar, between 3 g/L to 7 g/L, depending on the tested phage, of agar (Th. Geyer, Cat. # 214520) was added to LB medium before autoclaving. Growth medium was supplemented with 50 µg/mL kanamycin (Carl Roth, Cat. # T832. 3) or 100 µg/mL carbenicillin (Carl Roth, Cat. # 6344. 3) when appropriate. To induce expression from arabinose inducible promoters, 0. 5% L-arabinose (Sigma-Aldrich, A3256-25G) was added. To repress expression, 0. 5% glucose was used. After transformation, *E. coli* were recovered in SOC medium (20 g/L Tryptone, 5 g/L Yeast Extract, 0. 5 g/L NaCl, 2. 5 mM KCl, 10 mM MgCl2, 20 mM glucose, pH = 7. 0) at 37°C, shaking orbitally at 220 rpm.

### Phage cultivation and plaque assays

Appropriate *E. coli* host strains were grown in 3 mL of LB medium until OD_600_ = 0. 3 was reached. The bacterial culture was infected with either 10 µL of phage lysate or with a small amount of cryopreserved phages. The infected cultures were incubated 10 min at 37°C and afterwards moved to a shaker continuing the incubation shaking at 220 rpm. After 3 h or after the culture had cleared, 1. 8 mL were transferred to a fresh tube and mixed with chloroform (ITW Reagents, Cat. # 131252. 1612) to 0. 5% total concentration. The culture was vortexed briefly and then centrifuged 1 min at 11, 000 xg. The upper phase was filtered through a 0. 2 µm filter (Sarstedt, Cat. # 83. 1826. 001) and stored at 4°C until further use. For plaque assays the double agar overlay method was used. Individual colonies of host bacteria were used to inoculate 3 mL LB medium and were incubated 16h. The cultures were back diluted to an OD_600_ equal to 0. 05 and incubated until they reached OD_600_ = 0. 5. 2 mL of bacterial suspension was pelleted at 5, 000 xg for 3 min and resuspended in 200 µL of LB medium. 100 µL of this bacterial suspension was mixed with 3 mL pre-warmed soft agar, vortexed and poured on top of LB agar plates. The tested phages were serial diluted in SM buffer (5. 8 g/L NaCl, 2 g/L MgSO_4_, 50 mM Tris-HCl pH 7. 4) and 3 µL droplets were spotted on the solidified soft-agar overlay. The plates were incubated 16 h at 37°C and the formed plaques were used to enumerate PFU/mL.

### Plasmid construction

Plasmids were constructed using the NEBuilder HiFi DNA assembly (NEB, E2621L) according to manufacturer’s instructions. Unless otherwise specified, primers were designed with 30 - 50 nt complementary overhangs. For smaller mutations the NEB Q5 site-directed mutagenesis kit (NEB, Cat. # E0554S) was used according to the manufacturer’s protocol. For expression of *thsB1* under an arabinose-inducible promoter and *thsAB2* under their native promoter, plasmid pOD328 was constructed, incorporating mutations in the 5′ untranslated region that reduced background toxicity of the construct in the heterologous host.

### *E. coli* transformation

To make *E. coli* strains competent for electroporation, individual colonies were inoculated in 3 mL LB medium and incubated for 16 h. Afterwards, they were back diluted in 25 mL of fresh LB medium to an OD_600_ equal to 0. 05. The cultures were incubated until they reached an OD_600_ of 0. 5. The cultures were centrifuged at 5, 000 xg, and 4°C for 10 min and the supernatant was discarded. The pellet was washed with 10 mL of pre-cooled 10% glycerol (VWR, Cat. # J61059. AP) and centrifuged at 5, 000 xg and 4°C for 10 min. The supernatant was discarded, the cell pellet was resuspended in 1 mL cold 10% glycerol and centrifuged 1 min at 4°C and 11, 000 xg. The cell pellet was resuspended in 250 µL cold 10% glycerol and 50 µL aliquots were prepared per electroporation reaction. For the transformation of plasmid DNA, 50 ng of DNA was mixed with the electrocompetent cells and mixed carefully by pipetting. The suspension was transferred to a 0. 1 cm electroporation cuvette and a pulse of 1. 8 kV, 25 μF and 200 Ω was applied using the Gene Pulser Xcell Total Electroporation System (BioRad, Cat. # 1652660). The bacteria were recovered with 500 µL of SOC medium for 1 h at 37°C, before plating on LB plates containing the appropriate antibiotics.

### Infection dynamics in liquid culture

Host cultures were prepared from individual colonies inoculated in 3 mL of LB medium and incubated for 16 h. The next day they were back diluted in 180 µL of fresh medium aliquoted in a 96-well plate (Fisher Scientific, Cat. # 10212811) to an OD_600_ = 0. 05. 20 µl of phage lysate was added to the bacterial cultures to achieve the desired MOIs. The plate was incubated 10 min at 37°C without shaking, to allow the phages to adsorb to the bacteria. The 96-well plate was loaded in a BioTek Synergy H1 multimode microplate reader and incubated at 37°C while shaking at 425 rpm. OD_600_ measurements were taken in 5-minute intervals over 16 h in total.

### Reporter dye assays

Host cultures were prepared from individual colonies inoculated in 3 mL of LB medium and incubated for 16 h. The next morning they were back diluted in 180 µl of fresh LB medium, supplemented with 0. 5 mg/mL of BSA (PAN Biotech, Cat. # P06-1391100) and aliquoted in a 96-well plate to an OD_600_ equal to 0. 05. 20 µl of phage lysate was added to the bacterial cultures to achieve the desired MOIs. The cultures were incubated for 1 h at 37°C shaking at 425 rpm before adding the corresponding reporter dye. Resazurin (Sigma-Aldrich, Cat. # R7017-1G) was added to a final concentration of 3 µg/mL and fluorescence measured at 560 nm excitation / 590 nm emission. DiBAC4(3) (Biomol, Cat. # ABD-21411) was added to a final concentration of 5 µM and measured at 490 nm excitation / 516 nm emission. Propidium iodide (Life Technologies, Cat. # P3566) was added to a final concentration of 20 µM and measured at 537 nm excitation / 618 nm emission. The 96-well plate was loaded in a BioTek Synergy H1 multimode microplate reader and incubated at 37 °C while shaking at 425 rpm. Measurements were taken in 5 minute intervals for a total 16 h.

### Genomic modification of ATCC25922

To knock out the *thsAB2B1* locus from ATCC 25922, Lambda Red recombineering was used (50). First, ATCC 25922 was transformed with the pKD46 helper plasmid carrying the Lambda Red recombineering system. Transformation was performed by electroporation, as described above. Recovery and incubation were carried out at 30°C, since the plasmid is heat-sensitive. Individual colonies were inoculated into 3 mL of LB medium and grown for 16 h at 30°C with orbital shaking at 220 rpm. The cultures were then back-diluted by adding 100 µL into 25 mL LB medium supplemented with 0. 5% L-arabinose and incubated under the same conditions until reaching an OD_600_ of 0. 5. At this point, cells were prepared to be electrocompetent, as described above. A linear repair template was generated by PCR amplification of the FRT-flanked kanamycin resistance cassette from plasmid pKD13, using primers with 35-bp homology arms. The first and the last nucleotides of the homology regions correspond to positions 3, 089, 464 and 3, 086, 855, respectively, the *E. coli* ATCC 25922 genome (NCBI accession CP009072) **(Table S2)**. 1 µg of linear repair template was electroporated, and the bacteria were recovered in 1 mL SOC medium at 37°C with orbital shaking at 220 rpm for 4 h. The recovered cells were centrifuged at 5000 ×g for 3 min, the supernatant discarded, and the pellet resuspended in 100 µL of LB medium. Ten microliters of the suspension were plated on LB agar with carbenicillin to confirm loss of the helper plasmid, while the remaining cells were plated on LB agar with kanamycin. Correct integration of the resistance cassette was verified by PCR and Sanger sequencing (Microsynth AG). Confirmed clones were subsequently made electrocompetent as described above and transformed with the pCP20 helper plasmid encoding FLP recombinase using electroporation. Recovery and incubation were performed at 30°C due to plasmid heat sensitivity. Colonies were restreaked on LB agar with carbenicillin, with kanamycin, and without antibiotics, followed by incubation at 37°C for 16 h. Clones sensitive to both carbenicillin and kanamycin were further analyzed by Sanger sequencing to confirm correct excision of the resistance cassette.

### One-step growth curves

One-step growth curves were performed to evaluate the propagation of phages in their respective hosts. Host cultures were prepared from individual colonies inoculated in 3 mL LB medium and incubated for 16 h. The culture was back diluted in 3 mL of fresh LB medium and incubated until it reached an OD_600_ equal to 0. 3. The bacterial cultures were infected at MOI=1 with the corresponding phage. The culture was incubated at 37°C, shaking orbitally at 220 rpm and 200 µL samples were taken in 20 min intervals, for a total of 120 min. Phages were isolated from these cultures as described above. To enumerate the phages, plaque spot assays were performed on a susceptible host.

### NAD concentration assay

To evaluate intracellular NAD concentration of bacterial cultures, the NAD/NADH Assay Kit (Sigma-Aldrich, MAK468) was used. For the assay, 200 µl of bacterial culture was used. The assay was performed according to the manufacturer’s protocol. The NAD concentration was quantified by measuring the optical density at 565 nm over 15 min in a BioTek Synergy H1 multimode microplate reader and comparing it to the standard curve of purified NAD. To calculate the NAD concentration we used the formula:

NAD (µM) = (ΔOD_Sample_ - ΔOD_Blank_) / Slope (µM^-1^)

## ACKNOWLEDGMENTS

This work was supported through funding by the Deutsche Forschungsgemeinschaft (BE 6703/2-1 to C.L.B.), the European Research Council (865973 to C.L.B.), and the PostDoc Plus Program from the Graduate School of Life Sciences at Julius-Maximilians-Universität Würzburg (to O.D.).

## AUTHOR CONTRIBUTIONS

Conceptualization, writing–original draft, and visualization: all authors. System identification: O.D. Experiments in *E. coli*: F.E. and O.D. Structure modeling: F.E. Supervision and funding acquisition: C.L.B.

## MATERIAL AND DATA AVAILABILITY

All constructs and raw data can be shared upon reasonable request.

## Supplementary Information

### SUPPLEMENTARY FIGURES

**Figure S1:**
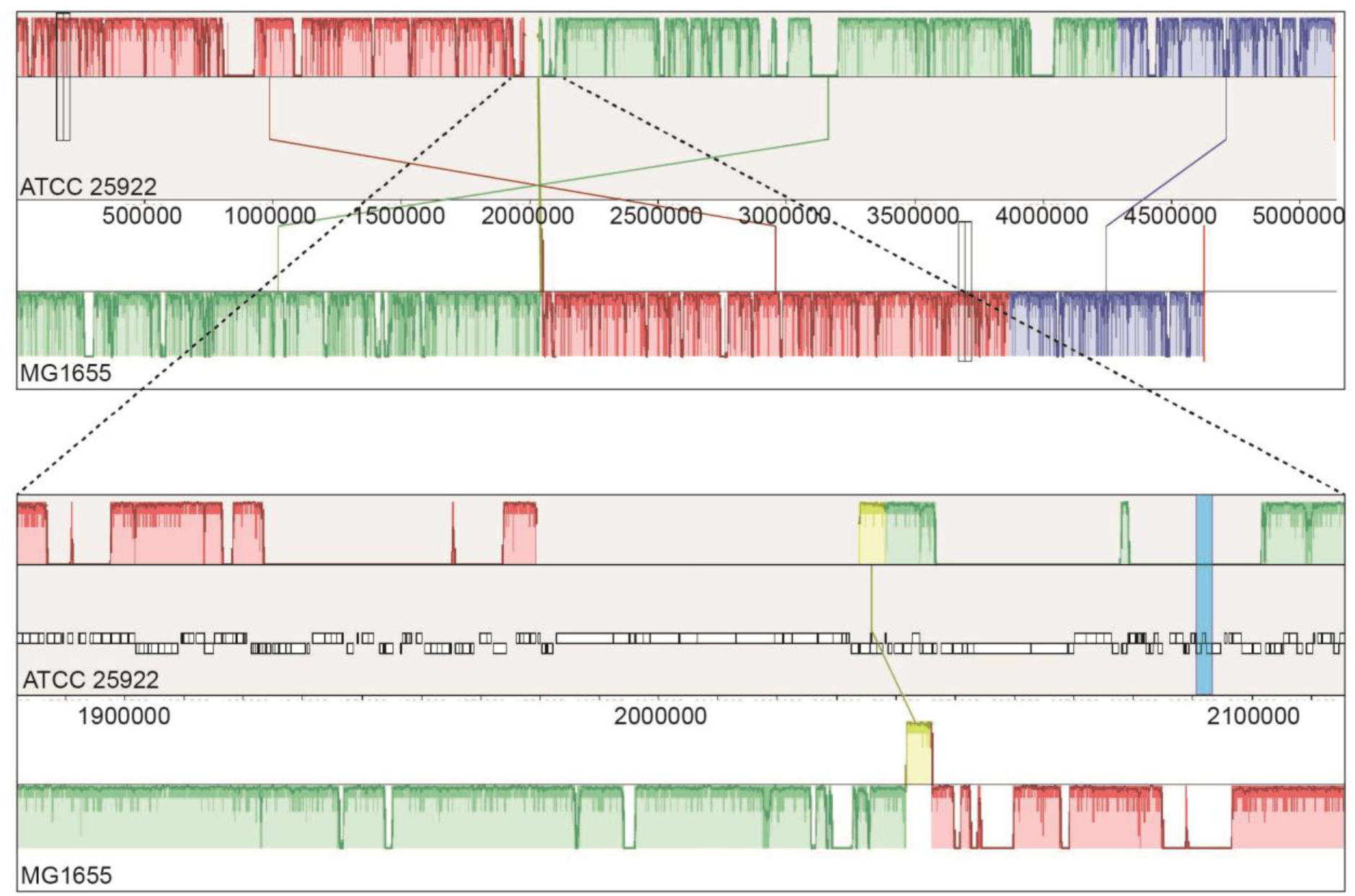
progressiveMAUVE alignment of ATCC 25922 and MG1655 genome sequences. The bottom shows a zoomed in view of the ATCC 25922 genomic region (1, 850, 000 - 2, 150, 000) containing the Thoeris locus, which is highlighted in blue. The height of the bar describes the similarity profile that corresponds to the average conservation between regions, while an empty space indicates a lack of detectable homology. The color of the bars show large regions that are presumably homologous and internally free from genomic rearrangement.

**Figure S2:**
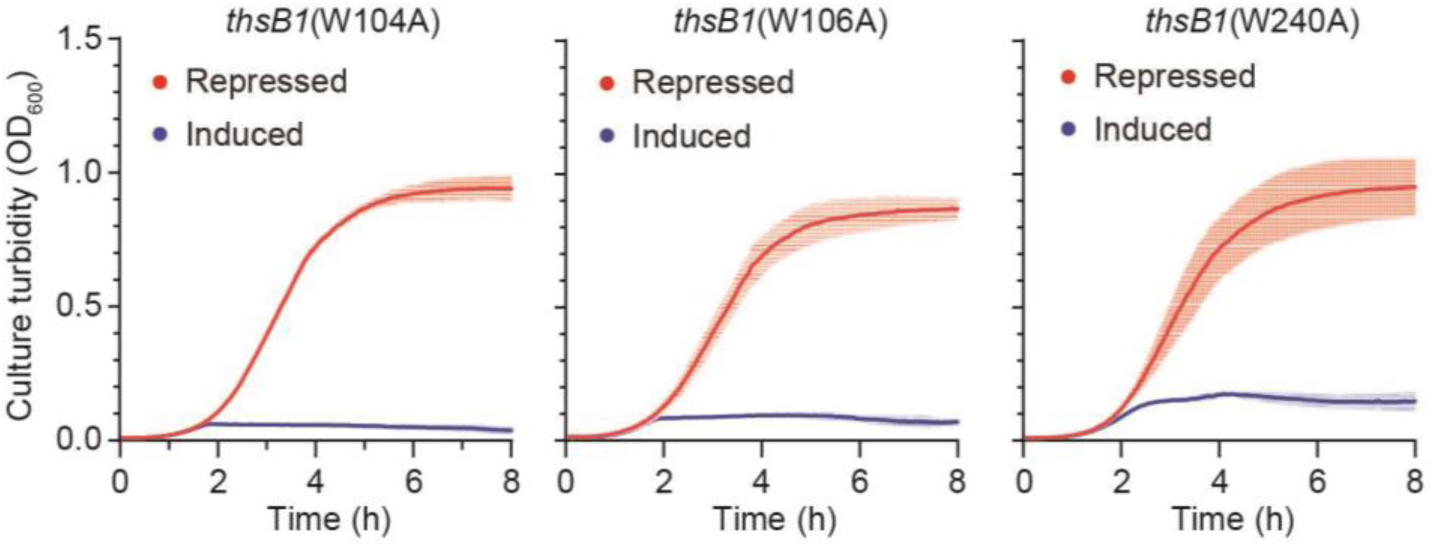
Growth curves of *E. coli* MG1655 heterologously expressing the *thsB1B2A* locus, with *thsB1* under the control of an arabinose inducible promoter. Every *thsB1* tryptophan residue that is part of a WxxxE motif has been individually mutated to alanine. *E. coli* have been cultured in either inducing (+ L-arabinose) or repressing (+ D-glucose) conditions. Each curve and shaded region represent the average and standard deviation of three independent cultures.

**Supplementary figure S3:**
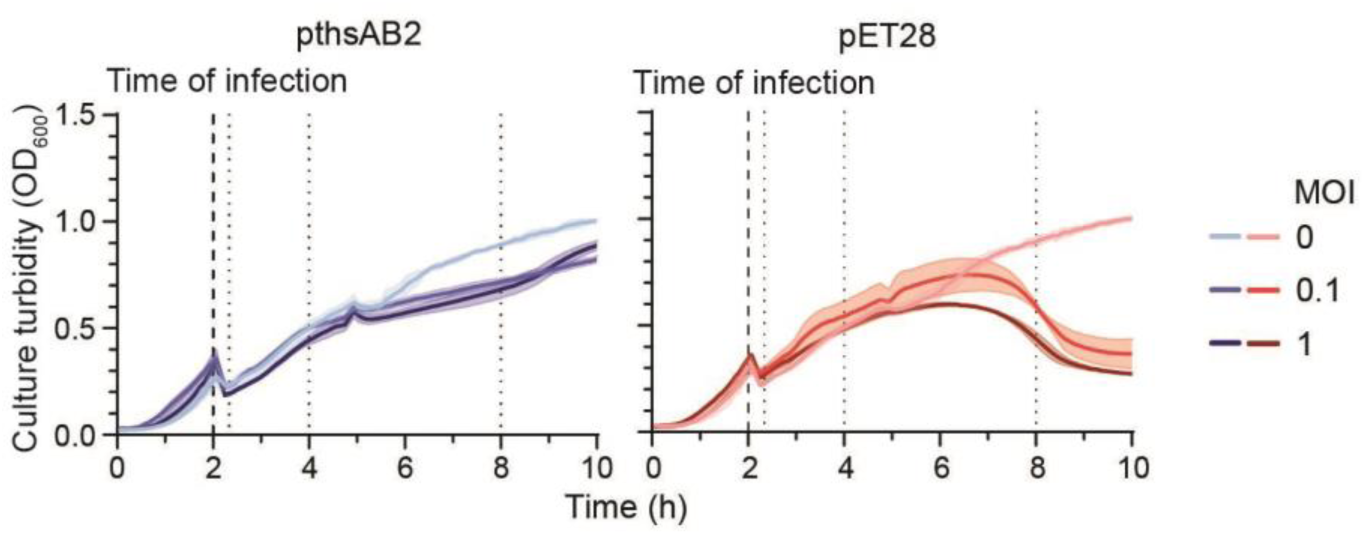
Growth curves of *E. coli* heterologously expressing *thsAB2* (pthsAB2) or an empty vector control (pET28). The bacteria were infected after 2 hours of incubation (dashed line), with phage T4 at different MOIs. Cultures were harvested for the NAD assay determining cellular NAD concentration 30 minutes, 2 hours and 6 hours after infection (dotted lines). Each curve shows the average of three independent replicates, with the shaded area indicating the standard deviation.

### SUPPLEMENTARY TABLES

**Table S1:** TM-align scores of ThsB1 and ThsB2 aligned with representative structures of the 11 Thoeris subfamilies. See the included Supplementary Table .xlsx file.

**Table S2:** Bacterial strains used in this study. See the included Supplementary Tables .xlsx file.

**Table S3:** Bacteriophages used in this study. See the included Supplementary Tables .xlsx file.

**Table S4:** Plasmids used in this study. See the included Supplementary Tables .xlsx file.

**Table S5:** Primers used in this study. See the included Supplementary Tables .xlsx file.

